# CRISPR-Cpf1 mediates efficient homology-directed repair and temperature-controlled genome editing

**DOI:** 10.1101/156125

**Authors:** Miguel A. Moreno-Mateos, Juan P. Fernandez, Romain Rouet, Maura A. Lane, Charles E. Vejnar, Emily Mis, Mustafa K. Khokha, Jennifer A. Doudna, Antonio J. Giraldez

## Abstract

Cpf1 is a novel class of CRISPR-Cas DNA endonucleases, with a wide range of activity across different eukaryotic systems. Yet, the underlying determinants of this variability are poorly understood. Here, we demonstrate that LbCpf1, but not AsCpf1, ribonucleoprotein complexes allow efficient mutagenesis in zebrafish and *Xenopus*. We show that temperature modulates Cpf1 activity by controlling its ability to access genomic DNA. This effect is stronger on AsCpf1, explaining its lower efficiency in ectothermic organisms. We capitalize on this property to show that temporal control of the temperature allows post-translational modulation of Cpf1-mediated genome editing. Finally, we determine that LbCpf1 significantly increases homology-directed repair in zebrafish, improving current approaches for targeted DNA integration in the genome. Together, we provide a molecular understanding of Cpf1 activity *in vivo* and establish Cpf1 as an efficient and inducible genome engineering tool across ectothermic species.

---

Cpf1 is a newly discovered class 2/type V CRISPR-Cas DNA endonuclease^1^ that displays a range of activity across different systems. Thus far, two Cpf1 proteins have been used for genome editing in mammalian cells: AsCpf1 and LbCpf1, which are derived from *Acidaminococcus sp BV3L6* and *Lachnospiraceae bacterium ND2006*, respectively^1^. CRISPR-Cpf1 presents several advantages for genome engineering, including i) extended target recognition in T-rich sequences (PAM 5′TTTN), such as non-coding RNAs, 5’ and 3’ UTRs (Supplementary Fig. 1a, Supplementary Table 1) ii) high specificity in mammalian cells^2,3^ and iii) shorter crRNA (~43nt), facilitating *in vitro* synthesis (Supplementary Fig. 1b-e). CRISPR-Cpf1 has been efficiently used to generate targeted mutations in mice^4-6^, however, AsCpf1 shows lower activity in *Drosophila* and plants^7-10^, hindering the broad application of Cpf1 across different model systems^7-10^. Here, we investigate the underlying determinants of these differences, and whether these insights can be exploited to optimize this method across species and modulate the mutagenic activity of Cpf1 *in vivo*.

In this study, we characterize and optimize the CRISPR-Cpf1 genome editing system in zebrafish (*Danio rerio*) and *Xenopus tropicalis*. We demonstrate that recombinant Cpf1 protein allows Cpf1-mediated genome editing. In contrast, delayed protein expression from mRNA-encoded Cpf1 results in rapid degradation of unprotected crRNAs *in vivo*. We show that temperature influences Cpf1 activity *in vivo,* affecting AsCpf1 more dramatically, which explains its lower activity in ectothermic organisms such as zebrafish, *Xenopus*, *Drosophila*^9^, and plants^7,8,10^. We capitalize on these differences to develop a method that provides temporal control of Cpf1-mediated mutagenesis, resulting in different onset and size mutant clones. Finally, we show that CRISPR-LbCpf1 together with a single stranded DNA (ssDNA) donor significantly increases the efficiency of homology-directed repair in zebrafish when compared to CRISPRCas9. All together, these results provide a highly efficient and inducible genome engineering system in ectothermic organisms.

## LbCpf1-crRNA RNP complexes provide a robust genome editing system in zebrafish and *X. tropicalis.*

To implement Cpf1-mediated genome editing we compared the activity of recombinant proteins and mRNAs encoding codon-optimized AsCpf1 or LbCpf1^1^ in zebrafish. We used these with three crRNAs targeting *slc45a2 (albino*) or *tyr* (*tyrosinase*), which are involved in pigmentation. We observed that ribonucleoprotein (RNP) complexes showed a dramatic increase in activity compared to mRNA delivery of Cpf1 (Fig. 1). While AsCpf1 and LbCpf1 mRNAs were efficiently translated and the proteins were detected by Western blot analysis (Supplementary Fig. 2a), different crRNAs were rapidly degraded after mRNA injection (Supplementary Fig. 2b,c). Precursor-crRNAs (pre-crRNAs)^11^ did not increase mutagenic activity nor the stability of the crRNA (Supplementary Fig. 2d-f). Interestingly, we observed a significant increase in crRNA half-life (Supplementary Fig 3a,b) when Cpf1-crRNA RNP complexes were preassembled before injection, suggesting that Cpf1 protein protects crRNAs from rapid degradation *in vivo.*

**Figure 1.**
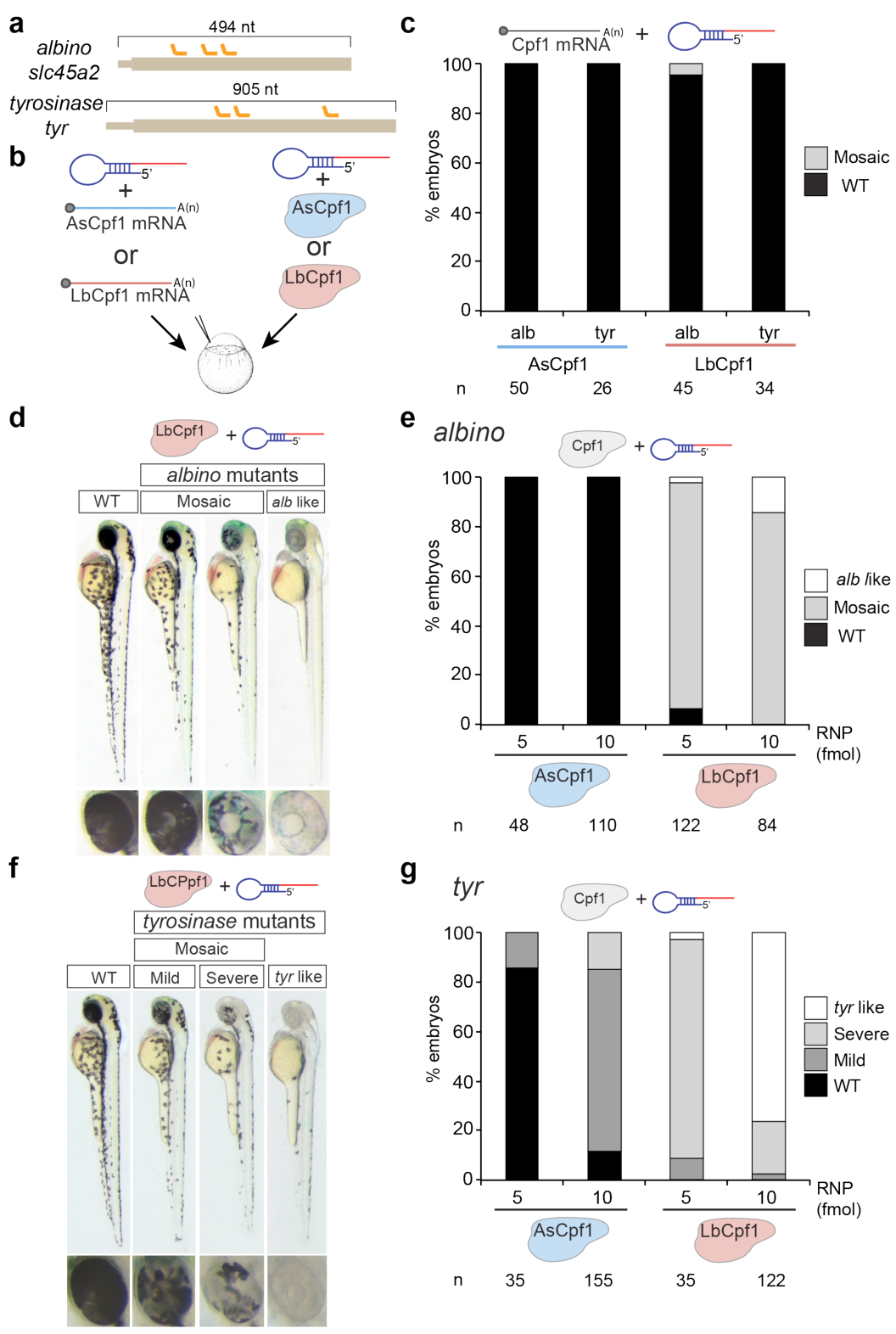
LbCpf1-crRNA RNP complexes are a robust genome editing system in zebrafish. **a.** Diagram illustrating 3 crRNAs (orange) targeting *slc45a2* and *tyr* exon 1 in zebrafish. **b.** Schematic illustrating the experimental setup to analyze CRISPR-Cpf1-mediated mutations in zebrafish. 3 crRNAs (panel a) were either mixed with mRNA coding for AsCpf1 or LbCpf1 or assembled into RNP complexes with their corresponding purified proteins and injected in one-cell stage embryos. **c.** Phenotypic evaluation of crRNAs (30 pg/ crRNA) and mRNA (100 pg) injections. Stacked barplots showing the percentage of mosaic (gray) and phenotypically wild-type (WT) (black) embryos 48 hours post-fertilization (hpf) after injection. **d.** Phenotypes obtained after the injection of the LbCpf1-crRNA RNP complexes targeting *slc45a2* showing different levels of mosaicism compared to the WT. Lateral views and insets of the eyes of 48 hpf embryos are shown. **e.** Phenotypic evaluation of Cpf1-crRNA RNP complexes injections targeting *slc45a2* (*albino*). Stacked barplots showing the percentage of *alb*-like (white), mosaic (gray), and phenotypically WT (black) embryos 48 hpf after injection using different amounts (fmol) of RNP complexes. Number of embryos evaluated (n) is shown for each condition. **f.** Phenotypes obtained after the injection of the LbCpf1-crRNA RNP complexes targeting *tyr* showing different levels of mosaicism compared to the wild type (WT). Lateral views and insets of the eyes of 48 hpf embryos are shown. **g.** Phenotypic evaluation of LbCpf1-crRNA RNP complexes targeting *tyrosinase* (*tyr).* Stacked barplots showing the percentage of *tyr-*like (white), severe mutant (light gray), mild mutant (dark grey) and phenotypically WT (black) embryos 48 hpf after injection using different amounts (fmol) of RNP complexes. Number of embryos evaluated (n) is shown for each condition.

Analysis of AsCpf1 and LbCpf1 RNP complexes revealed that both cleaved DNA *in vitro*. However, only LbCpf1 induced efficient mutagenesis in zebrafish (Fig. 1d-g, Supplementary Fig. 3c-f, Supplementary Data 1). In contrast, AsCpf1 showed very low mutagenic activity (Fig. 1e,g). Similar results were obtained when targeting the same loci in *Xenopus tropicalis* (Supplementary Fig. 4, Supplementary Data 1). To estimate the rate of germ-line transmission for each locus, we quantified the number of *albino* or *tyrosinase* homozygous loss-of-function mutants in the offspring of the zebrafish embryos injected with LbCpf1-crRNA RNP complexes. We obtained ~88% and ~99% for *alb* and *tyr* respectively (Supplementary Fig. 3g,h), demonstrating a high level of mutagenesis in the germ cells. Collectively, these results demonstrate that, in contrast to LbCpf1 mRNA injection, delivery of pre-assembled LbCpf1-crRNA RNP complexes provides a robust genome editing tool in zebrafish and *X. tropicalis*.

## Temperature modulates Cpf1 activity *in vitro* and *in vivo*

While AsCpf1 efficiently functions *in vitro* and in mammalian cells at 37°C^1,4-6^, its activity is dramatically reduced in zebrafish, *X. tropicalis, Drosophila*^9^, and plants^7,8,10^ which develop below 28°C. Thus, we hypothesized that temperature may impact AsCpf1 activity *in vivo*. Consistently, we observed that AsCpf1 is less active than LbCpf1 at 25°C and 28°C, although both proteins show comparable cleavage activity at 37°C (Supplementary Fig. 4d, Supplementary Fig. 5a).

To determine whether temperature modulates Cpf1 activity *in vivo* we compared the mutagenic activity of AsCpf1- and LbCpf1-injected embryos raised at different temperatures. We observed that a brief incubation of embryos at 34°C post-injection significantly increased AsCpf1-mediated gene editing (Fig. 2a-c), measured by an increase in the number of mosaic mutant embryos and the severity of the phenotype (Fig. 2b,c, Supplementary Fig. 5b). Similarly, LbCpf1 activity increased at higher temperatures, generating ~70% *albino*-like and ~100% *tyrosinase*-like mutants (Fig. 2d,e, Supplementary Fig. 5b,c). Higher temperatures also increased Cpf1 activity in *X. tropicalis* (Supplementary Fig. 5d), indicating that this effect is observed across different ectothermic organisms. In contrast to Cpf1, higher temperatures did not modulate the activity of SpCas9-sgRNA RNP complexes *in vivo* or *in vitro* (Supplementary Fig. 6). All together, these results suggest that temperature specifically influences Cpf1 function *in vitro* and *in vivo*. This effect is stronger in AsCpf1, thus explaining the lower efficiency of this protein in ectothermic organisms.

**Figure 2.**
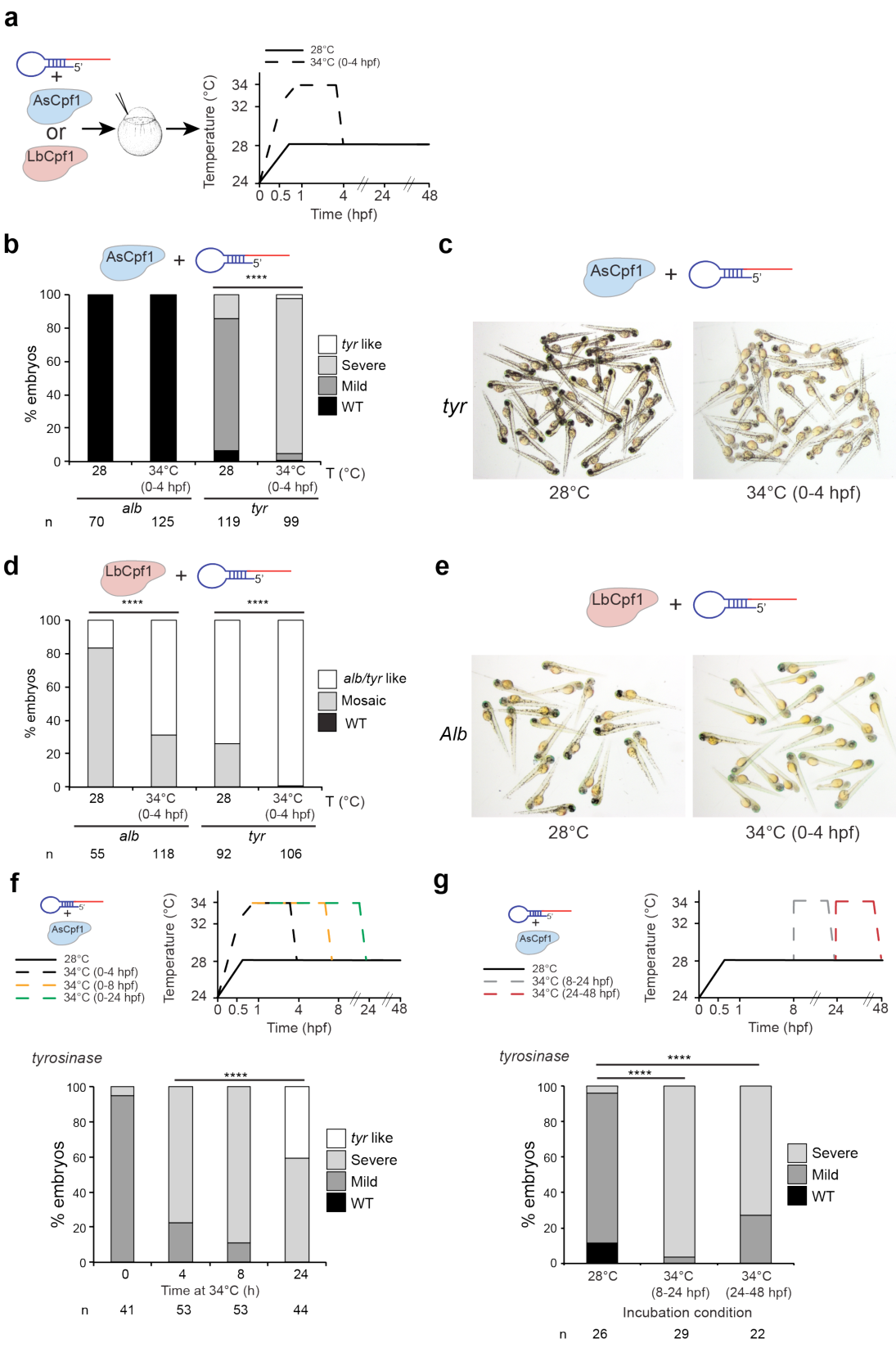
Temperature is a key factor modulating Cpf1 activity *in vitro* and *in vivo*. **a.** Schema illustrating different temperature incubations after Cpf1-crRNA RNP complexes injections targeting *slc45a2* (*alb*) and *tyr.* **b.** Phenotypic evaluation of AsCpf1-crRNA RNP complexes (10 fmol) injections at different temperature incubations (T). Stacked barplots showing the percentage of *tyr*-like (white), severe mutant (light gray), mild mutant (dark grey) and phenotypically WT (black) embryos 48 hpf after injection. Number of embryos evaluated (n) is shown for each condition. χ^2^ test (**** p< 0.0001). **c.** A representative picture showing 48 hpf old embryos obtained after AsCpf1 - crRNA RNP complexes injections targeting *tyr* at different temperature incubations. d. Phenotypic evaluation of LbCpf1-crRNA RNP complexes (10 fmol) injections at different temperature incubations (T). Stacked barplots showing the percentage of *alb*/*tyr-*like (white), mosaic mutants (grey) and phenotypically WT (black) embryos 48 hpf after injection. Number of embryos evaluated (n) is shown for each condition. χ^2^ test (**** p< 0.0001). **e.** A representative picture showing 48 hpf old embryos obtained after crRNA-LbCpf1-crRNA RNP complexes injections targeting *slc45a2* at different temperature incubations. **f.** Schematic illustrating different incubation conditions (0,4,8 or 24 h at 34°C, then 28°C) after crRNA-AsCpf1 RNP complexes (10 fmol) injections targeting *tyr* in zebrafish (top). Phenotypic evaluation of crRNAs-AsCpf1 RNP complexes injections targeting *tyr* in the conditions described above (bottom). Stacked barplots showing the percentage of *tyr-*like (white), severe mutant (light gray), mild mutant (dark grey), and phenotypically WT (black) embryos 48 hpf after injection. Number of embryos evaluated (n) is shown for each condition. χ^2^ test (****p< 0.0001). **g.** Schematic illustrating different incubation conditions: 8h at 28°C, 16 h at 34°C and then 24 h at 28°C (34°C 8-24 hpf) or 24 h at 28°C, then 24 h at 34°C (34°C 24-48 hpf) after crRNA-AsCpf1 RNP complexes (10 fmol) injections targeting *tyr* in zebrafish (top). Phenotypic evaluation of crRNAs-AsCpf1 RNP complexes injections targeting *tyr* in the conditions described above (bottom). Stacked barplots showing the percentage of severe mutant (light gray), mild mutant (dark grey), and phenotypically WT (black) embryos 48 hpf after injection. Number of embryos evaluated (n) is shown for each condition. χ^2^ test (****p< 0.0001).

Next, we asked whether temperature control of AsCpf1 activity could be used to modulate genome editing over time. First, we observed that longer incubations at 34°C increased the number of mutant cells and further improved AsCpf1 efficiency *in vivo* to a level that was comparable to LbCpf1 at 28°C (Fig. 2f, Supplementary Fig. 7). This suggests that the crRNA-Cpf1 complex is still active and able to mutagenize the genome later in development, allowing for potential on-off modulation of the mutagenic activity with temperature over time. To test this possibility, we compared the mutagenic activity of AsCpf1 at 28°C to embryos incubated at 34°C at 8-24 hpf or 24-48 hpf. In both cases, we observed an increase in the severity of the phenotype and the extent of mutant cells, suggesting that modulating AsCpf1 activity over time can control the onset of mutagenesis and the number of independent mutant events generated in vivo (Fig. 2g). Together, these results suggest that temperature sensitivity of AsCpf1 activity can be used to modulate genome editing over time.

Based on the differential activity of Cpf1, we hypothesized that temperature may control Cpf1 endonuclease activity and/or accessibility to the genomic DNA. Previous studies have shown that targeting catalytically dead SpCas9 (SpdCas9) to a genomic locus improves accessibility to the flanking regions of the DNA. This method, called proxy-CRISPR technique, restores the activity of inactive Cpf1 and Cas9 orthologues in mammalian cells^12^. To understand the molecular basis of the temperature modulation of Cpf1 activity, we delivered a proxyCRISPR adjacent (29-32 nt) to each Cpf1 targeted loci. We observed that proxy-CRISPR enhanced LbCpf1 (Fig. 3a,d,e), and AsCpf1 activity (Fig. 3a-c) at 28°C, suggesting that temperature likely influences the competence of Cpf1 to access or unwind genomic DNA rather than its endonuclease activity.

**Figure 3.**
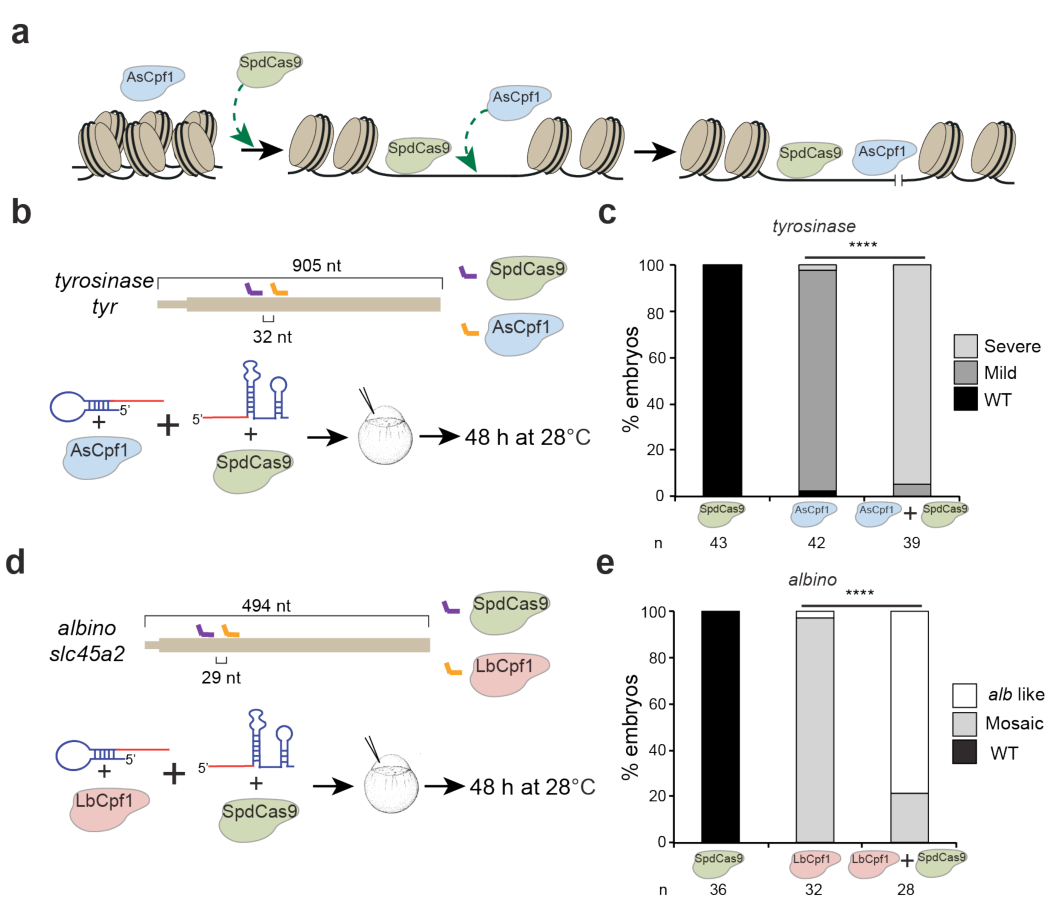
Catalytic dead SpCas9 (SpdCas9) proximal targeting increases Cpf1 activity. **a.** Schematic diagram of proxy-CRISPR approach. Temperature may control Cpf1 activity to access and/or unwind genomic DNA, impeding AsCpf1 to access the genomic target *in vivo* at 28°C. When SpdCas9 binds in the proximity of the AsCpf1 target, it facilitates the availability of AsCpf1 to cleave the inaccessible target. **b.** Diagram illustrating crRNA 1 (orange) and a proximal sgRNA (purple) targeting *tyr* exon 1 (Supplementary Table 1) in zebrafish (top). AsCpf1-crRNA and/or SpdCas9-sgRNA RNP complexes were injected into one-cell stage embryos and then incubated at 28°C for 48 h (bottom). **c.** Phenotypic evaluation of the experiment described in b. Stacked barplots showing the percentage of severe mutant (light gray), mild mutant (dark grey), and phenotypically WT (black) embryos 48 hpf after injection. Number of embryos evaluated (n) is shown for each condition. χ^2^ test (**** p< 0.0001). **d.** Diagram illustrating crRNA 2 (orange) and sgRNA 2 (purple) targeting *alb* exon 1 (Supplementary Table 1, Supplementary Fig. 3f, Supplementary Fig. 6e) in zebrafish (top). LbCpf1-crRNA and/or SpdCas9-sgRNA RNP complexes were injected into one-cell stage embryos and then incubated at 28°C for 48 h (bottom) **e.** Phenotypic evaluation of the experiment described in d. Stacked barplots showing the percentage of *alb*-like (white), mosaic mutants (grey) and phenotypically WT (black) embryos 48 hpf after injection. Number of embryos evaluated (n) is shown for each condition. χ^2^ test (**** p< 0.0001).

## LbCpf1 enhances homology-directed repair (HDR) compared to SpCas9 in zebrafish

Cas9-mediated HDR is low and highly variable in zebrafish^13-15^. One potential drawback of SpCas9 is that SpCas9-induced indels can prevent recurrent cleavage of the target DNA by generating base-pairing mismatches between the “seed” sequence of the target and the sgRNA (Supplementary Fig. 1e)^16^. In contrast, Cpf1 induces a double-strand break (DSB) 18 nt away from the PAM sequence (Supplementary Fig. 1c). Thus we hypothesized that repeated cleavages without destroying the target site may increase the window of opportunity to repair DSB through HDR. Encouraged by the high activity of LbCpf1 in zebrafish, we tested the capability of this endonuclease to facilitate HDR-mediated DNA integration in zebrafish and compared it to SpCas9 across two different loci (Fig. 4, Supplementary Fig. 8). To this end, we used 10 different ssDNA donors, with a similar structure to those previously tested to optimize Cas9-mediated HDR in cell culture^17^. The main features tested included single-strand DNA donors that were i) centered on the 3’end of the DSB, ii) complementary to either the target or the non-target strand (which contains the PAM sequence) and iii) with different homology arm lengths (Fig. 4a-c, Supplementary Fig. 8d-f). We observed that LbCpf 1 increases HDR up to ~4 fold (as average of percentage of HDR per embryo) when compared to optimized ssDNA donor for SpCas9 (Fig. 4d, Supplementary Fig. 8g)^17^. In contrast to SpCas9, Cpf1 causes the highest HDR rate when the single strand donor DNA is complementary to the target strand (Fig. 4d, Supplementary Fig. 8g). Altogether, these results suggest that LbCpf1 activity in combination with ssDNA donors complementary to the target strand strongly enhances HDR in zebrafish.

**Figure 4.**
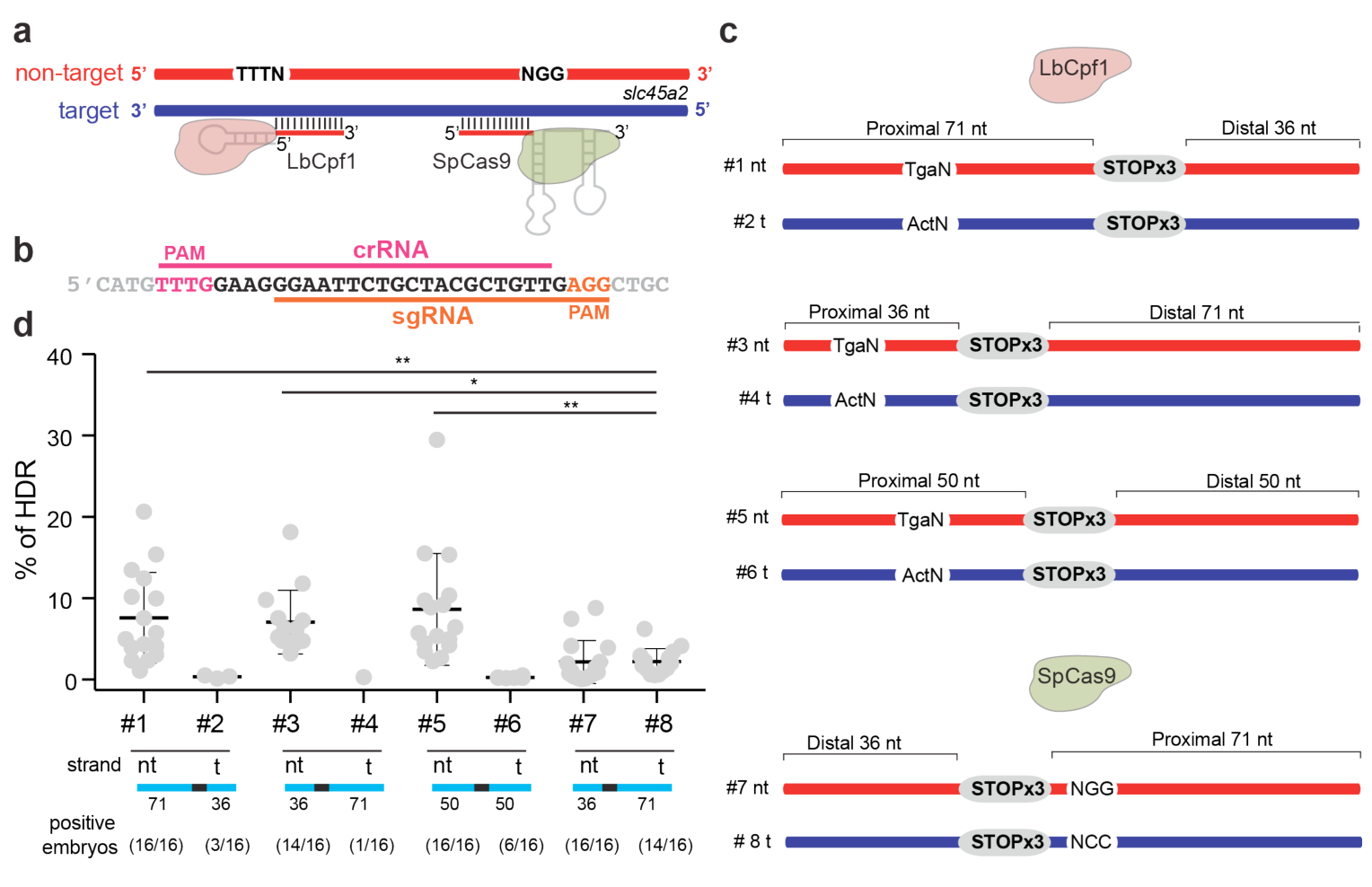
LbCpf1-mediated homology directed repair. **a.** Schematic illustrating LbCpf1-crRNA RNP complexes and sgRNA SpCas9-sgRNA RNP complexes interaction with their respective DNA targets. crRNA and sgRNA-DNA annealing occurs on the target strand (blue), and PAM sequences are in the non-target strand (red). **b.** crRNA (pink line) and sgRNA (orange line) overlapping target sequences in the *albino* locus used for this analysis. **c.** Schema illustrating different donor ssDNA (#1-#6) complementary to either the target strand (t) or non-target strand (nt) and with symmetric or asymmetric homology arms used in combination with LbCpf1-crRNA. PAM sequence was modified (TgaN/ActN) to prevent new editing post-HDR. An optimized ssDNA donor (#8) described for SpCas9-induced HDR^17^ and its complementary version (#7) were used in combination with SpCas9-sgRNA RNP complexes as references for comparison (bottom). **d.** qPCR quantification showing % of HDR from individual embryos when using LbCpf1 and different ssDNA donors in comparison to SpCas9. Results are shown as the averages ± standard errors of the means from 16 embryos in two independent experiments (n=8 embryos per experiment). Embryos showing a PCR amplification signal detectable by qPCR were considered positive. The data were analyzed by one-way ANOVA, followed by Bonferroni post-test for significance versus control condition (#8), (*) P < 0.05, (**) P < 0.01.

## Discussion

In this study, we optimize the CRISPR-Cpf1 system to generate mutations in non-mammalian vertebrates. In particular, we demonstrate that i) LbCpf1 RNP complexes efficiently mutagenize the genomes of zebrafish and *X. tropicalis*; ii) AsCpf1 activity is lower at 28°C and regulated by temperature, providing a method to modulate mutagenic activity over time and iii) in the absence of Cpf1 protein, crRNAs are unstable and rapidly degraded *in vivo,* explaining the lack of activity when Cpf1 mRNA is injected in zebrafish^6^. Interestingly, previous studies have shown reduced activity of AsCpf1 in *Drosophila*^9^ and across various plant species^7,8,10^. Our findings provide a context to understand how AsCpf1 activity is reduced in ectothermic organisms. We demonstrate that the lower activity of AsCpf1 at 28°C is recovered when SpdCas9 facilitates the genomic DNA accessibility or when zebrafish embryos are incubated at 34°C. This temperature control can be useful to address later mutant phenotypes for genes that function at different developmental stages. Overall, our results suggest that LbCpf1 is a more suitable as a constitutive nuclease in ectothermic animals, while AsCpf1 allows temperature and temporal control of the nuclease activity during development.

Moreover, we demonstrate that LbCpf1 achieves higher HDR rates than SpCas9 in zebrafish, likely by allowing repeated cleavages before indel mutations terminate targeting^1^, supported by the longer deletions caused by Cpf1 compared to Cas9^2,18^ (Supplementary Data 1). Repeated cleavages would increase the window of opportunity to repair DSB through HDR instead of alternative end joining, which is the main repair pathway during zebrafish development^15^. Interestingly, our results indicate that ssDNA donors complementary to the target sequence dramatically increase LbCpf1-mediated HDR. This is in contrast to SpCas9, where previous studies have shown that the highest HDR efficiency is achieved with asymmetric ssDNA donors complementary to the non-target strand^17^. These results suggest that Cpf1 may first release the PAM-proximal target strand, becoming available for HDR. Overall, our results provide insights into the molecular mechanisms used by distinct endonucleases to release one DNA strand and enable subsequent repair *in vivo*.

Finally, we have developed an online resource tool to predict all potential Cpf1 targets and off-targets across *C. elegans*, sea urchin, sea anemone, *Drosophila*, zebrafish, medaka, *Xenopus*, chicken, mouse, rat, and human genomes in an updated, publicly-available resource CRISPRscan: may2017.crisprscan.org. Altogether, this study will guide optimization strategies for the CRISPR-Cpf1 system across ectothermic organisms.

## Acknowledgements

We thank H. Codore and K. Bishop for technical help; J-D Beaudoin, V. Tornini and C. Fellmann for discussions and all the members of the Giraldez laboratory for intellectual and technical support; M. Fernandez-Fuertes for helping with western blot analysis and C. Takacs and V. Tornini for manuscript editing. Programa de Movilidad en Áreas de Investigación priorizadas por la Consejería de Igualdad, Salud y Políticas Sociales de la Junta de Andalucía (M.A.M-M.), NIH grants R21 HD073768, R01 HD074078, GM103789, GM102251, GM101108 and GM081602 (A.J.G.), The Swiss National Science Foundation grant P2GEP3_148600 (C.E.V) and R01 HD081379, 4R33HL120783 (E.M., M.A.L. and M.K.K.) supported this work. M.K.K. is supported by the Edward Mallinckrodt Jr Foundation. A.J.G is supported by the HHMI Faculty Scholar program, the March of Dimes, the Yale Scholars Program and the Whitman fellowship funds provided by E.E. Just, L.B. Lemann, E. Evelyn and M. Spiegel, the H. Keffer Hartline and E.F. MacNichol Jr at the Marine Biological Laboratory in Woods Hole, MA. M.K.K. is supported by the Edward Mallinckrodt Jr Foundation. R.R. acknowledges support from the Australian National Health and Medical Research Council for his early career postdoctoral fellowship (APP1090875). J.A.D. is an investigator of the Howard Hughes Medical Institute.

## Author contributions

M.A.M-M. and A.J.G. conceived the project and M.A.M-M., R.R., and A.J.G. designed the research. M.A.M-M. performed zebrafish experiments and *in vitro* assays, R.R. purified and provided recombinant proteins, J.P.F. designed and performed HDR experiments with help from M.A.M-M. E.M. and M.A.L. carried out *Xenopus* injections and phenotype analyses. C.E.V determined target and off target sites for different CRISPR systems, developed crisprscan.org and help with statistical analysis. M.A.M-M., R.R., J.P.F., A.J.G. and J.A.D. performed data analysis and M.A.M-M., R.R. and A.J.G. wrote the manuscript with input from the other authors. M.K.K. provided reagents and materials. All authors reviewed and approved the manuscript.

## Competing interests

The authors declare no competing financial interests related to this work. J.A.D. is employed by the Howard Hughes Medical Institute (HHMI) and works at the University at California (UC), Berkeley, USA. UC Berkeley and HHMI have patents pending for CRISPR technologies on which J.A.D. is an inventor. J.A.D. is the executive director of the Innovative Genomics Institute (IGI) at UC Berkeley and University of California, San Francisco. J.A.D. is a co-founder of Editas Medicine, Intellia Therapeutics and Caribou Biosciences, and a scientific adviser to Caribou, Intellia, eFFECTOR Therapeutics and Driver.

## Methods

### crRNA and sgRNA target sites design

Target sites were designed using an updated version of CRISPRscan (may2017.crisprscan.org) tool^18^. sgRNAs (5′GGN19GG) and crRNAs (5′TTTN24) target sites without predicted off-targets were used^19,20^.

### crRNA, sgRNA and Cpf1 mRNA generation

crRNA and pre-crRNA DNA template were generated by fill-in PCR (Supplementary Fig. 1d and Supplementary Fig. 2d). A crRNA or pre-crRNA (As/Lb) universal primer (Supplementary Table 1) containing the T7 promoter (5’-TAATACGACTCACTATA-3), and the mature crRNA repeat or the complete pre-crRNA repeat for AsCpf1 or LbCpf1 preceded by 5’GG were used in combination with a specific oligo of 43 nt adding the spacer (target binding sequence) and the repeat sequence (reverse complement orientation). A 65/66 (crRNA) or 80/ 81 (pre-crRNA) bp PCR product was generated following these conditions: 3 minutes at 95°C, 30 cycles of 30 seconds at 95°C, 30 seconds at 52°C and 20 seconds at 72°C, and a final step at 72°C for seven minutes. PCR products were purified using Qiaquick PCR purification kit (Qiagen) columns and used as template (200-250 ng) for a T7 *In vitro* transcription (IVT) reaction (AmpliScribe-T7-Flash transcription kit from Epicentre; 6h of reaction). *In vitro* transcribed crRNAs were DNAse treated and precipitated with Sodium Acetate/Ethanol. crRNAs were visualized in a 2% agarose stained by ethidium bromide to check for RNA integrity. sgRNAs were generated as previously described^18,21^. sgRNAs and crRNAs targeting *slc45a2 in* zebrafish were individually *in vitro* transcribed. Pre-crRNAs targeting *slc45a2* in zebrafish and crRNAs targeting *slc45a2* and *tyr* in *X. tropicalis* were in vitro transcribed from a pool of PCR templates for each crRNA per gene. Solid phase extraction purified crRNA targeting *tyr* in zebrafish were purchased (Synthego).

A zebrafish codon-optimized AsCpf1 and a human codon optimized LbCpf1^1^ were PCR amplified using the following primers: 5’-TTTTccATGGGCACCCAGTTCGAGGGA-3’ and 5’-TTTTCCGCggTTATCCGGCGTAATCGGGCACGTC3 ‘for Ascpf1 and 5’-ttttGCGGCCGCCACCATGAGCAAGCTGGAGAAGT T-3’ and 5’ ttttgaattcTTAGGCATAGTCGGGGACAT-3’ and 5’ ttttgaattcTTAGGCATAGTCGGGGACAT-3’ for LbCpf1. The following PCR products were then digested with *NcoI* and *SacII* (AsCpf1) or *NotI* and *EcoRI* (LbCpf1), and ligated into the pT3TS-nCas9n^22^ and pSP64T plasmids previously digested with these enzymes, respectively. Final constructs were confirmed by sequencing. For making AsCpf1 or LbCpf1 mRNA, the template DNA was linearized using *XbaI* and mRNA was synthetized using the mMachine T3 or SP6 kit (Ambion), respectively. *In vitro* transcribed mRNAs were DNAse treated and purified using the RNeasy Mini Kit (Qiagen).

### Protein expression and purification

*E.coli* codon-optimized SpCas9, SpdCas9 AsCpf1 and LbCpf1 were cloned into pET-based bacterial expression plasmid. Proteins (with N-terminal 6xHis and MBP tags and C-terminal 2xSV40 tags) were expressed in *E. coli* Rosetta 2 in TB media at 16°C for 18 h following induction with 0.4 mM IPTG. Cells were lysed in 20 mM HEPES pH 7.5, 500 mM KCl, 20 mM imidazole, 5 mM TCEP, 10% glycerol (supplemented with protease inhibitors) by sonication. Proteins in the lysate were first captured onto Ni-NTA resin (Qiagen), washed with lysis buffer, and eluted with 20 mM HEPES pH 7.5, 100 mM KCl, 300 mM imidazole, 5 mM TCEP, 10% glycerol. 6xHis-MBP tag was removed by TEV protease cleavage. Proteins were next dialyzed to 20 mM HEPES pH 7.5, 100 mM KCl, 5 mM TCEP, 10% glycerol and captured onto an ion-exchange column (HiTrap Heparin, GE Healthcare). Proteins were eluted with a linear gradient of 100 mM to 1 M KCl. Finally, size exclusion chromatography (Superdex 200, GE Healthcare) was performed in 20 mM HEPES pH 7.5, 300 mM KCl for Cpf1 or 150 mM KCl for Cas9, 1 mM TCEP and 10% glycerol. Protein was concentrated and filtered, concentration measured by Abs_280 nm_ (Nanodrop, Thermo Fisher) and stored at -80°C.

### RNA and RNP injections

100 pg of *cpf1* mRNA and 30 pg of each crRNA/pre-crRNA were injected at the one-cell stage, respectively. crRNAs or sgRNA were resuspended in 20 mM HEPES pH 7.5, 1 mM TCEP, 1 mM MgCl_2_, 10% glycerol, 300 or 150 mM KCl, respectively, at 24 *μ*M and stored at -80°C. Cpf1-crRNA or Cas9-sgRNA RNP were prepared as follows: Cpf1 or Cas9 were diluted to 20 *μ*M in 20 mM HEPES pH 7.5, 1 mM TCEP, 1 mM MgCl_2_, 10% glycerol 300 mM or 150 mM KCl, respectively, and 10 *μ*L were added to 10 *μ*L of crRNA/pre-crRNA or sgRNA at 24 μM (Protein-RNA ratio 1:1.2). RNPs were incubated at 37°C 10 min and then kept at room temperature before use. RNPs were stored at -80°C and up to three cycles of freeze-thaw cycles maintained similar efficiency. One nL (10 fmol) or 0.5 nL (5 fmol) from 10*μ*M solution were injected at the one-cell stage zebrafish embryos. For HDR experiments 10 fmol of RNP complexes and 40 pg of ssDNA donor were injected in one-cell stage zebrafish embryos.

### Quantitative RT-PCR

For crRNA experiment, 0.5 nL of Cpf1 RNP complexes were injected at the one-cell stage and 10 embryos were collected at 0,2, and 5 hours post fertilization (hpf). Total RNA was isolated from embryos injected using TRIzol reagent (Life Technologies). 1 μg of purified total RNA was then subjected to reverse transcription using the SuperScript^®^ III First Strand Synthesis System (Thermo Fisher Scientific), using random hexamers and a specific primer for each crRNA (Supplementary Fig. 2b, Supplementary Table 1) following manufacturer’s protocol. 5 μL from a 1/50 dilution of the cDNA reaction was used to determine the levels of different crRNAs in a 20 μL reaction containing 1 μL of each oligo crRNA specific and universal primers or forward and reverse (10 μM) (Supplementary Table 1), using Power SYBR Green PCR Master Mix Kit (Applied Biosystems) and a ViiA 7 instrument (Applied Biosystems). PCR cycling profile consisted of incubation at 50 °C for 2 min, followed by a denaturing step at 95 °C for 10 min and 40 cycles at 95 °C for 15 s and 60 °C for 1 min. *taf15* and *cdk2ap2* genes expressed at similar levels during the first 5 hours postfertilization were used as normalization controls^23^.

For HDR experiments, 28 hpf injected zebrafish embryos were collected for DNA extraction^24^. Briefly, embryos were incubated in 80 μL of 100mM NaOH at 95 °C for 15 min. Next, 40 μL of Tris-HCl 1M pH 7.5 was added. Crude DNA extracts were 1/5 diluted (bi-distilled water) and 5 μL were used for qPCR with the corresponding forward and reverse primers (10 μM) (Supplementary Table 1, Supplementary Fig. 8), and using the same conditions described above. % HDR was measured using integration primers (a-c,f-i) and genomic DNA primers (d,e, j,k) (Supplementary Table 1, Fig 4, Supplementary Fig. 8). To calculate percentage of HDR, standard curves of different genomic qPCR products were performed using known amounts of DNA gBlocks^®^ gene fragments (Supplementary Table 1). Then, percentage of HDR per embryo was calculated dividing the amount of integrated DNA by total amount of genomic DNA.

### Western blot

Ten embryos were collected at 6 hpf and transferred to 200 μL of deyolking buffer for washing (55mM NaCl, 1.8 mM KCl, 1.25 mM NaHCO3). Deyolking buffer was discarded and 200 μL of the same buffer were added to resuspend the embryos by pipetting. The resuspended embryos were incubated at room temperature for 5 min with orbital shaking, then centrifuged at 300 x*g* for 30 s and washed with 110mM NaCl, 3.5 mM KCl, 10 mM Tris-HCl pH 7.4, 2.7 mM CaCl_2_. The pellet was resuspended in SDS sample buffer before separation by SDSPAGE and transfer to PVDF membrane. Anti-HA antibody (Roche) (1:1000) and rabbit polyclonal Anti-actin antibody (Sigma) (1:5000) were used according to manufacturer’s instructions. Secondary antibodies were fluorescence-labeled antibodies (alexa fluor 680) from LI-COR Biotechnology. Protein bands were visualized using the Odyssey Infrared Imaging System (LI-COR Biosciences, Lincoln, NE, USA).

### *In vitro* cleavage assays

The assays were carried as described in Jinek et al., 2012^25^ with minor modifications. Briefly, *slc45a2* and *tyr* targeted region (exon1) from zebrafish or *X. tropicalis* were amplified by PCR (Supplementary Table 1) and PCR products purified using QIAquick PCR purification kit (Qiagen). 100 ng of PCR purified product of *slc45a2* zebrafish (~ 0.11 pmol), *tyr* zebrafish (~ 0.23 pmol), *slc45a2 X. tropicalis* (~ 0.45 pmol), *tyr X. tropicalis* (~ 0.38 pmol) were subjected to in vitro cleavage with different concentrations of Cpf1 or SpCas9 RNP complexes in cleavage buffer (20 mM HEPES pH 7.5, 150 mM KCl, 0.5 mM DTT, 0.1 mM EDTA, 10mM MgCl2) at different temperatures for 90 min followed by 37°C for 5 min incubations with proteinase K (20 μg). The reactions were stopped with SDS loading buffer (30% glycerol, 0.6% SDS, 250 mM EDTA) and loading 1.5% agarose gel stained by ethidium bromide.

### Zebrafish maintenance, mating and image acquisition

Zebrafish wild-type embryos were obtained from natural mating of TU-AB and TLF strains of mixed ages (5–17 months). Selection of WT or LbCpf1-crRNA RNP in-jected mosaic F0 mutants mating pairs was random from a pool of 48 males and 48 females allocated for a given day of the month or random from the F0 mosaic mutants adult fish obtained ~3 months after injection, respectively. Fish lines were maintained in accordance with AAALAC research guidelines, under a protocol approved by Yale University IACUC.

All experiments were carried out at 28°C or 34°C, temperatures allowing normal development^26^.

Embryos were analyzed using a Zeiss Axioimager M1 and Discovery microscopes and photographed with a Zeiss Axiocam digital camera. Images were processed with Zeiss AxioVision 3.0.6.

### Frog husbandry and injections

*X. tropicalis* were housed and cared for in Khokha Lab aquatics facility according to established protocols approved by Yale IACUC.

*In vitro* fertilization and microinjection were done at 23.5°C, as previously described^27,28^. 2nL of RNP at 10 μM (20 fmol) were injected into one-cell stage embryo. After injection, embryos were left in 3% Ficoll for 30 minutes, then transferred to growing medium (1.1 mM Mg_2_Cl, 2.2 mM Ca_2_Cl, 2 mM KCl, 11mM NaCl, 5.5 mM HEPES pH 7.4) and incubated at different temperatures allowing normal development^28^.

### Statistics

Bar graphs are represented with s.d. error bars. Unpaired t-tests or one-way ANOVA were performed and P values were calculated with Prism (GraphPad Software, La Jolla, CA, USA).

---

**Supplementary figure 1.**
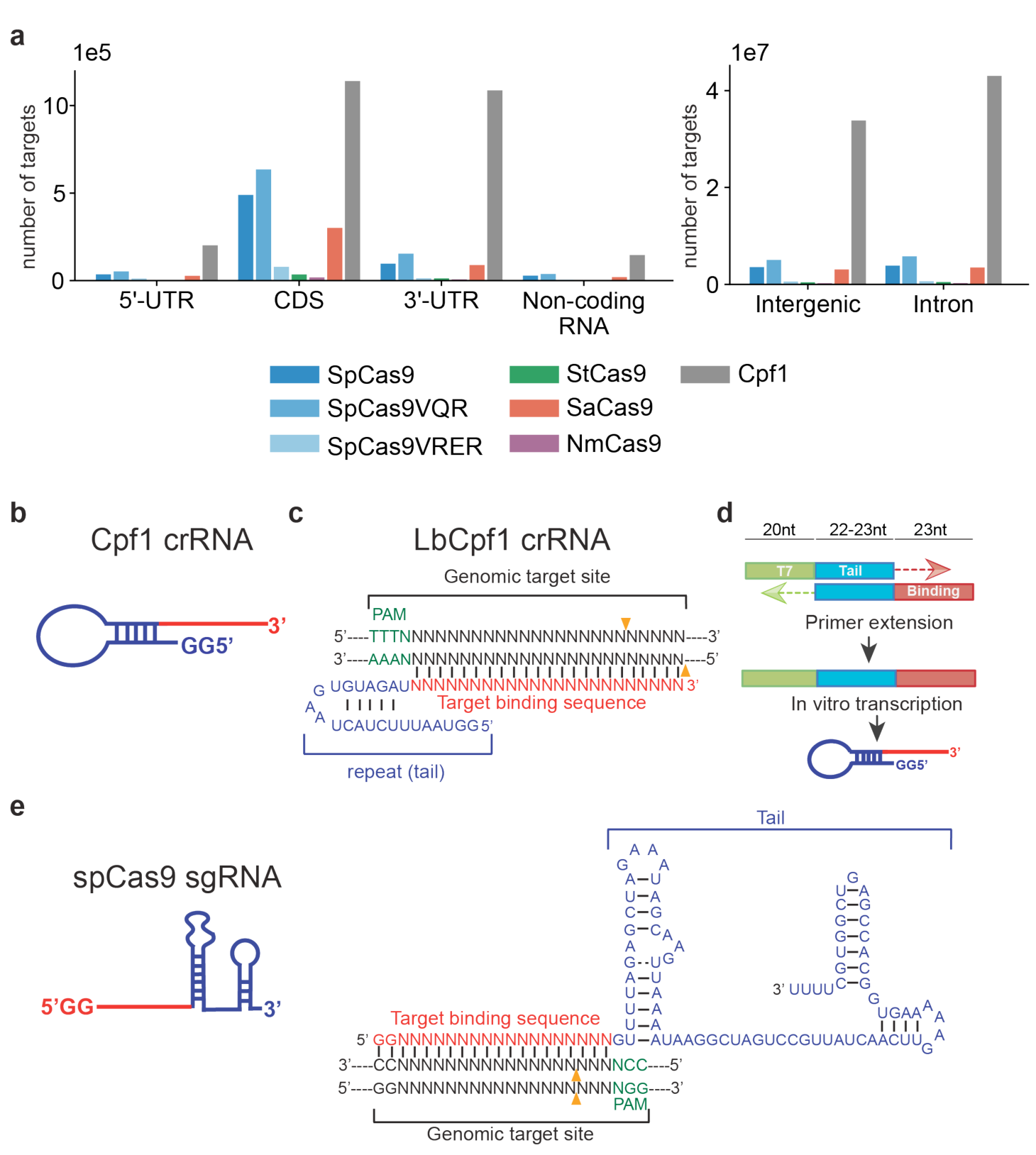
CRISPR-Cpf1 system features. **a.** Bar graphs showing number of targets of different CRISPR endonucleases in the zebrafish genome. Defined PAM sequences are available in Supplementary Table 1. SpCas9: *Streptococcus pyogenes* Cas9^1^; SpCas9VQR: *S. pyogenes* Cas9 VQR mutant^2^; SpCas9VRER: *S. pyogenes* Cas9 VRER mutant^2^; StCas9: *Streptococcus thermophilus* CRISPR 1 Cas9^3^; SaCas9: *Staphylococcus aureus* Cas9^4^; NmCas9: *Neisseria* meningiditis Cas9^3,5^; Cpf1: *Acidaminococcus sp BV3L6* / *Lachnospiraceae bacterium ND2006* Cpf1^6^. **b.** Schematic of crRNA structure. **c.** Scheme showing a LbCpf1 crRNA (binding sequence in red, repeat in blue) binding to the genomic target site (black) and the PAM sequence 5’-TTTN (green). Orange triangles indicate predicted cleavage sites. **d.** PCR approach to obtain a 65bp (AsCpf1) or 66bp (LbCpf1) bp product used as template for crRNA *in vitro* transcription. An oligonucleotide containing the T7 promoter (green) followed by two Guanine and 20 (AsCpf1) or 21 nt (LbCpf1) of the invariable repeat (tail, in blue) for annealing is used in combination with oligonucleotide containing the reverse complement of the repeat and 23 nt of the binding sequence (in red). **e.** Schematic of sgRNA structure (left) and scheme showing a sgRNA (binding sequence in red, tail in blue) binding to the genomic target site (black) and the PAM sequence 5’-NGG (green). Orange triangles indicate predicted cleavage sites. Adapted from Moreno-Mateos et al.^7^ While sgRNA is ~100nt, crRNA is ~43nt, which facilitates *in vitro* synthesis.

**Supplementary figure 2.**
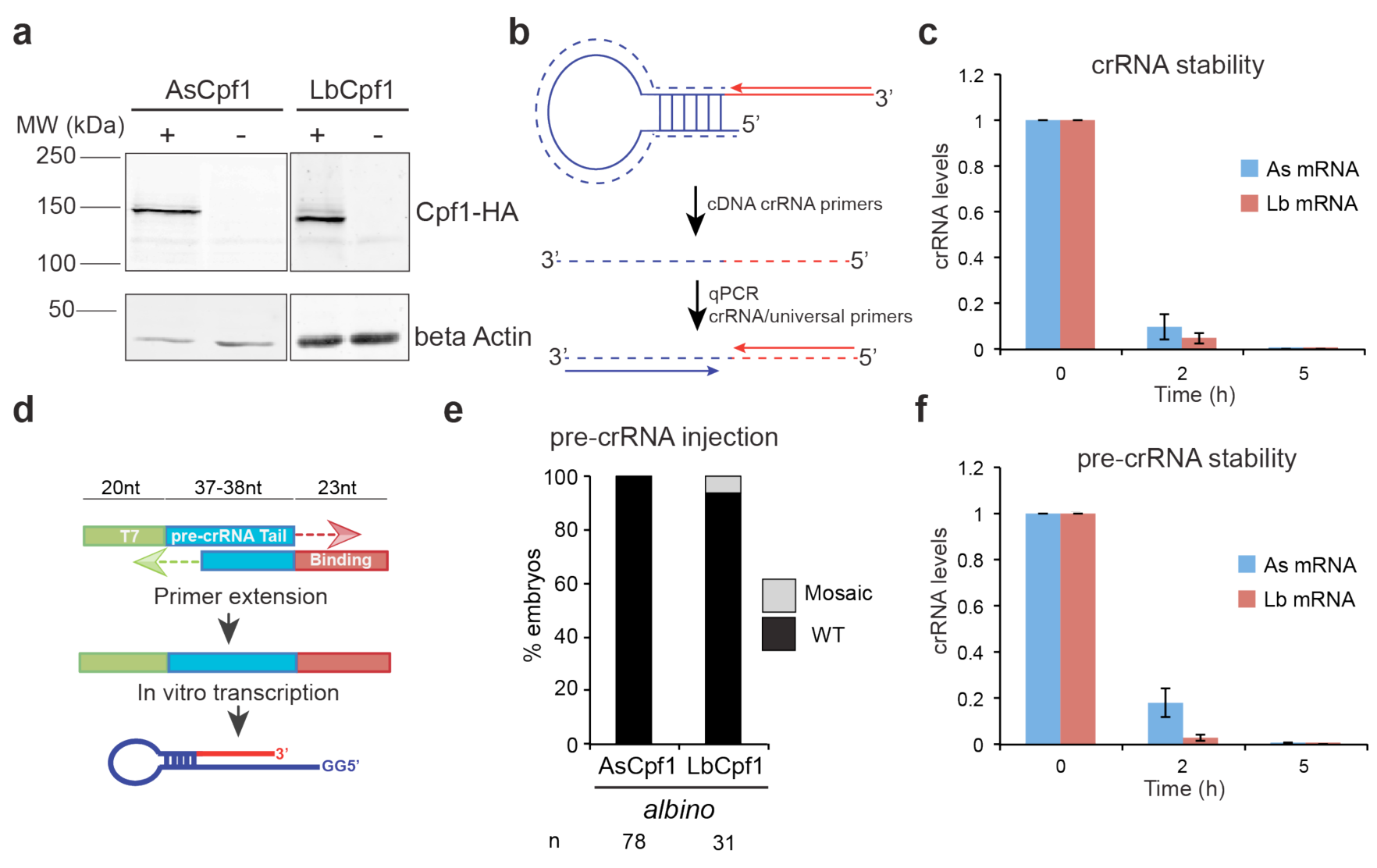
crRNA are rapidly degraded in zebrafish. **a.** Western blot showing Cpf1-HA protein in injected zebrafish embryos. Injected embryos (+) and uninjected (-) embryos as negative control were analyzed by using HA and beta-Actin antibodies. **b.** Schematic illustrating crRNA-cDNA and qPCR strategy. Specific crRNA primers annealing to the target binding sequence (red) were used to make cDNA. qPCR was carried out using a universal primer annealing to the repeat region (blue) and the specific primers described above (Supplementary Table 1). **c.** qRT–PCR analysis showing levels of crRNAs used for targeting *slc45a2* in Fig. 1c. Results are shown as the averages ± standard deviation of the mean for three crRNAs. **d.** PCR approach to obtain a 80bp (AsCpf1) or 81bp (LbCpf1) product used as template for pre-crRNA *in vitro* transcription similar to that described in Supplementary Fig. 1d, except the oligonucleotide containing the T7 promoter (green) includes the 35 nt (AsCpf1) or 36 nt (LbCpf1) pre-crRNA repeat sequence (tail). **e.** Phenotypic evaluation of pre-crRNAs (30 pg/pre-crRNA) and mRNA (100 pg) injections. Stacked barplots showing the percentage of mosaic (gray) and phenotypically WT (black) embryos 48 hpf after injection. **f.** qRT–PCR analysis showing levels of crRNAs used for targeting *slc45a2* in Supplementary Fig. 2e. Results are shown as the averages ± standard deviation of the mean for three crRNAs.

**Supplementary figure 3.**
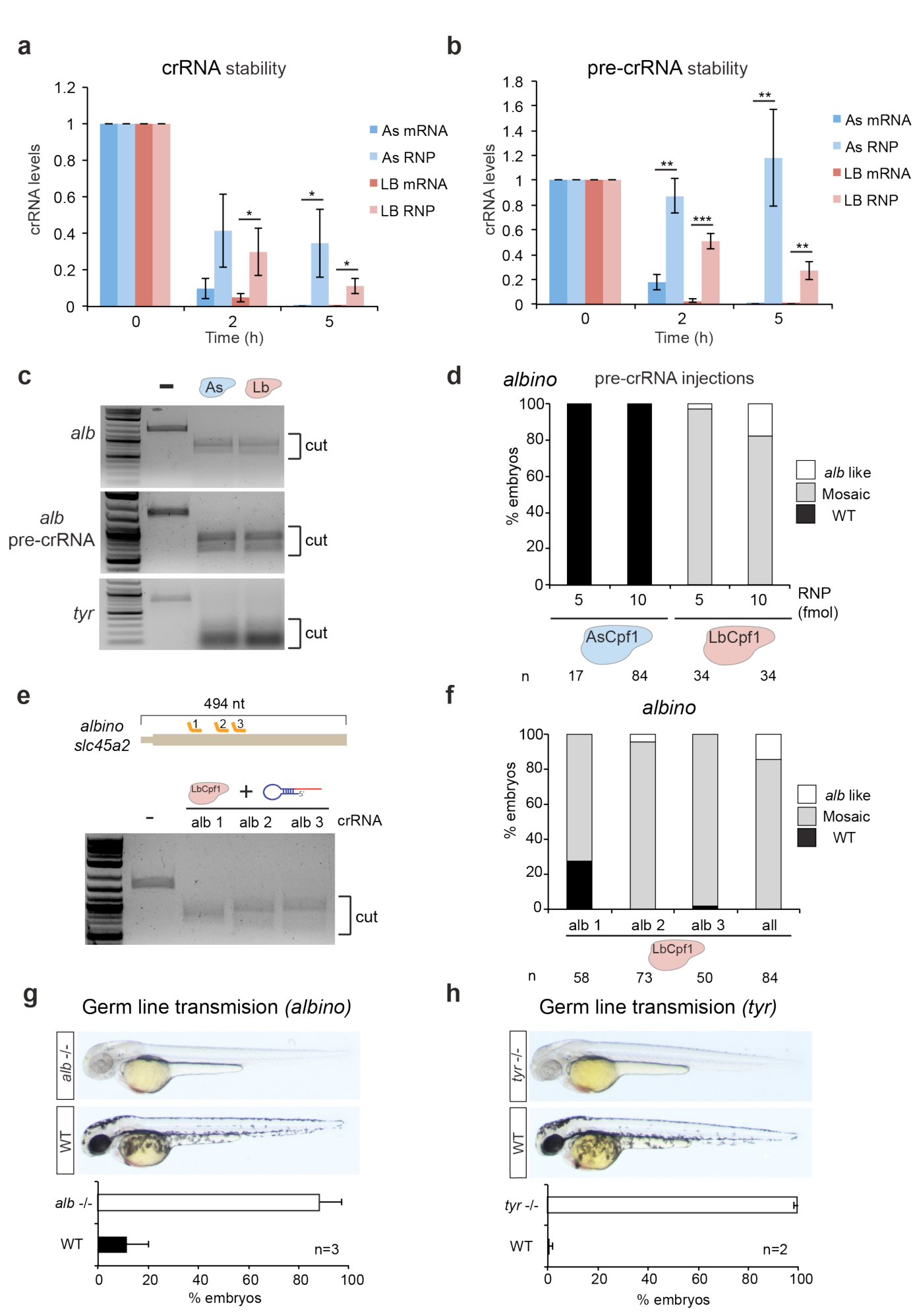
Cpf1-crRNA RNP complexes stabilize crRNA but only LbCpf1 allows robust genome editing in zebrafish. **a.** qRT–PCR analysis showing levels of crRNAs used for targeting *slc45a2* in Fig. 1c and 1e (5 fmol). Results are shown as the averages ± standard deviation of the mean for three crRNAs. The data were subjected to two-tailed Student’s t-test. (*) P< 0.05. **b.** qRT–PCR analysis showing levels of crRNAs used for targeting *slc45a2* in Supplementary Fig. 2e and 3d (5 fmol). Results are shown as the averages ± standard deviation of the mean for three crRNAs. The data were subjected to two-tailed Student’s t-test. (**) P < 0.01, (***) P < 0.001. **c**. In vitro cleavage assay using 10 pmol of AsCpf1 RNP or LbCpf1 RNP complexes containing a mix of 3 crRNAs (Fig. 1a) and a ~1.6 Kb (~ 0.11 pmol) PCR product including crRNA target sites for *slc45a2* (*alb*) or 20 pmol of AsCpf1 RNP or LbCpf1 RNP complexes containing a mix of 3 crRNAs and a 0.77 Kb (~ 0.23 pmol) PCR product including crRNA target sites for *tyrosinase*. Incubations were carried out at 37°C for 90 min. **d**. Phenotypic evaluation of precrRNAs-Cpf1 RNP complexes injections targeting *slc45a2*. Stacked barplots showing the percentage of *alb*-like (white), mosaic (gray) and phenotypically WT (black) embryos 48 hpf after injection. Number of embryos evaluated (n) is shown for each condition. **e.** In vitro cleavage assay using 10 pmol of LbCpf1 RNP complexes containing a individual crRNAs and a ~1.6 Kb (~ 0.11 pmol) PCR product including crRNA target sites for *slc45a2*. Incubations were carried out at 37°C for 90 min. **f.** Phenotypic evaluation of individual crRNAs-Cpf1 RNP (10 fmol) complexes injections targeting *slc45a2*. Stacked barplots showing the percentage of *alb*-like (white), mosaic (gray) and phenotypically WT (black) embryos 48 hpf after injection. Number of embryos evaluated (n) is shown for each condition. **g.** Phenotypes obtained from F0 *slc45a2* (*albino*) mutants in crosses (lateral views). Percentage of *alb* -/- and WT obtained from independent (n) in crosses. **h.** Phenotypes obtained from F0 *tyr*osinase mutants in crosses (lateral views). Percentage of *tyr* -/- and WT obtained from independent (n) in crosses.

**Supplementary figure 4.**
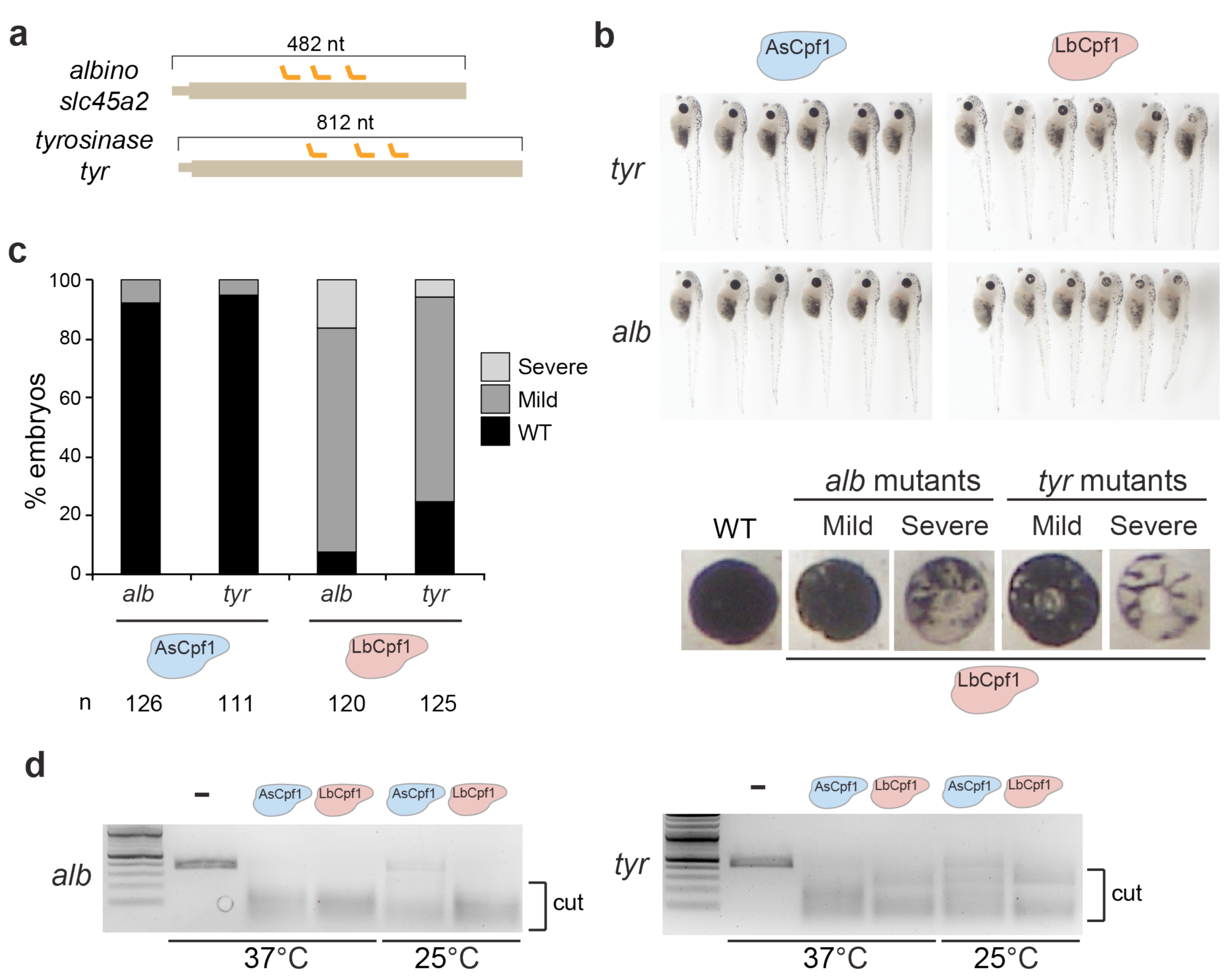
LbCpf1-crRNA RNP complexes are an efficient genome editing system in *X. tropicalis*. **a.** Diagram illustrating 3 crRNAs (orange) targeting *slc45a2* and *tyr* exon 1 in *X. tropicalis*. **b.** Phenotypes obtained after injection of the crRNA-AsCpf1 or crRNA-LbCpf1 RNP complexes (20 fmol) containing a mix of 3 crRNAs targeting *slc45a2* (*alb*) or *tyr* in *X. tropicalis*. Lateral views (top) and insets of the eyes (bottom) of stage 47–48 embryos are shown. **c.** Phenotypic evaluation of crRNAs-Cpf1 RNP complexes (20 fmol) injections targeting *slc45a2* (*alb*) or *tyr* in *X. tropicalis*. Stacked barplots showing the percentage of severe mutant (light gray), mild mutant (dark grey) and phenotypically WT (black) embryos at stage 47-48. Number of embryos evaluated (n) is shown for each condition. **d.** In vitro cleavage assay using 30 pmol of AsCpf1 RNP or LbCpf1 RNP complexes containing a mix of 3 crRNAs (Supplementary Fig. 4a) and a ~380pb (~ 0.45 pmol) or ~440pb (~ 0.38 pmol) PCR product including crRNA target sites for *slc45a2* (*alb*) (left) and *tyr* (right), respectively. Incubations were carried out at 37°C and 25°C for 90 min.

**Supplementary figure 5.**
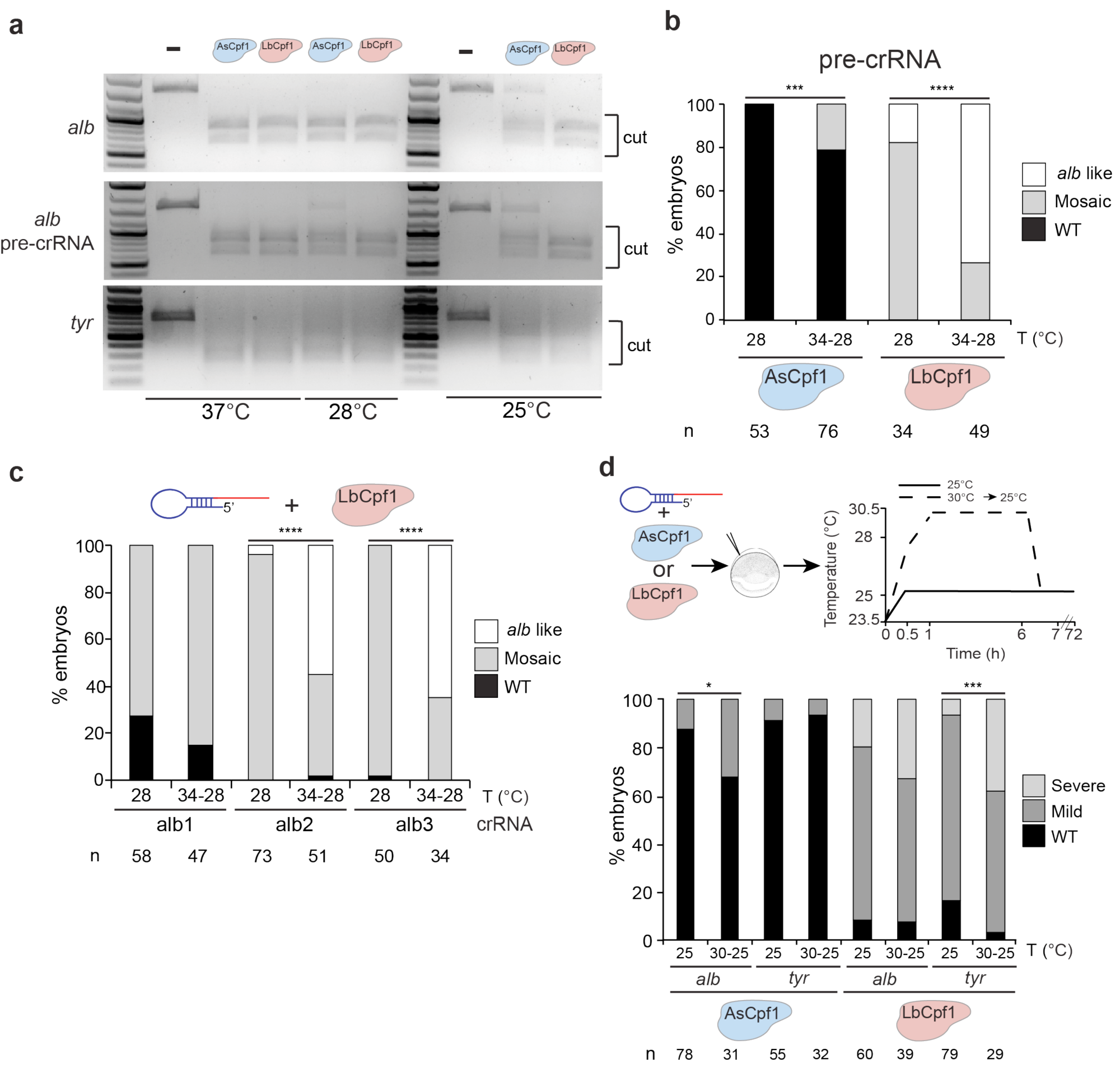
Temperature is a key factor modulating Cpf1 activity *in vitro* and *in vivo*. **a.** In vitro cleavage assay using AsCpf1 RNP or LbCpf1 RNP complexes containing a mix of 3 crRNAs (10 pmol) or pre-crRNAs (2.5 pmol) and a ~1.6 Kb (~ 0.11 pmol) PCR product (top and middle, respectively) containing crRNA target sites for *slc45a2* (*alb*) or a mix of 3 crRNAs (2.5 pmol) and a 0.77 Kb (~ 0.23 pmol) PCR product and containing crRNA target sites for *tyr*. Incubations were carried out at 37°C, 28°C or 25°C for 90 min. **b.** Phenotypic evaluation of pre-crRNAs-LbCpf1 RNP complexes (10 fmol) injections targeting *slc45a2* at different temperature incubations (T) (Fig. 2a). Stacked barplots showing the percentage of *alb*-like (white), mosaic mutants (grey) and phenotypically WT (black) embryos 48 hpf after injection. Number of embryos evaluated (n) is shown for each condition. χ^2^ test (*** p< 0.001,**** p< 0.0001). **c.** Phenotypic evaluation of individual crRNAs-LbCpf1 RNP complexes (10 fmol) injections targeting *slc45a2* at different temperature incubations (T) (Fig. 2a). Stacked barplots showing the percentage of *alb-*like (white), mosaic mutants (grey) and phenotypically WT (black) embryos 48 hpf after injection. Number of embryos evaluated (n) is shown for each condition. χ^2^ test (**** p< 0.0001). **d.** Schematic illustrating different temperature incubations after crRNA-Cpf1 RNP complexes injections targeting *slc45a2* (*alb*) and *tyr* in *X. tropicalis* (top). Phenotypic evaluation of pre-crRNAs-LbCpf1 RNP complexes (10 fmol) injections targeting *slc45a2 and tyr* at different temperature incubations (T) (bottom). Stacked barplots showing the percentage of severe mutant (light gray), mild mutant (dark grey) and phenotypically WT (black) embryos at stage 47-48. Number of embryos evaluated (n) is shown for each condition. χ^2^ test (*p< 0.05, ***p< 0.001).

**Supplementary figure 6.**
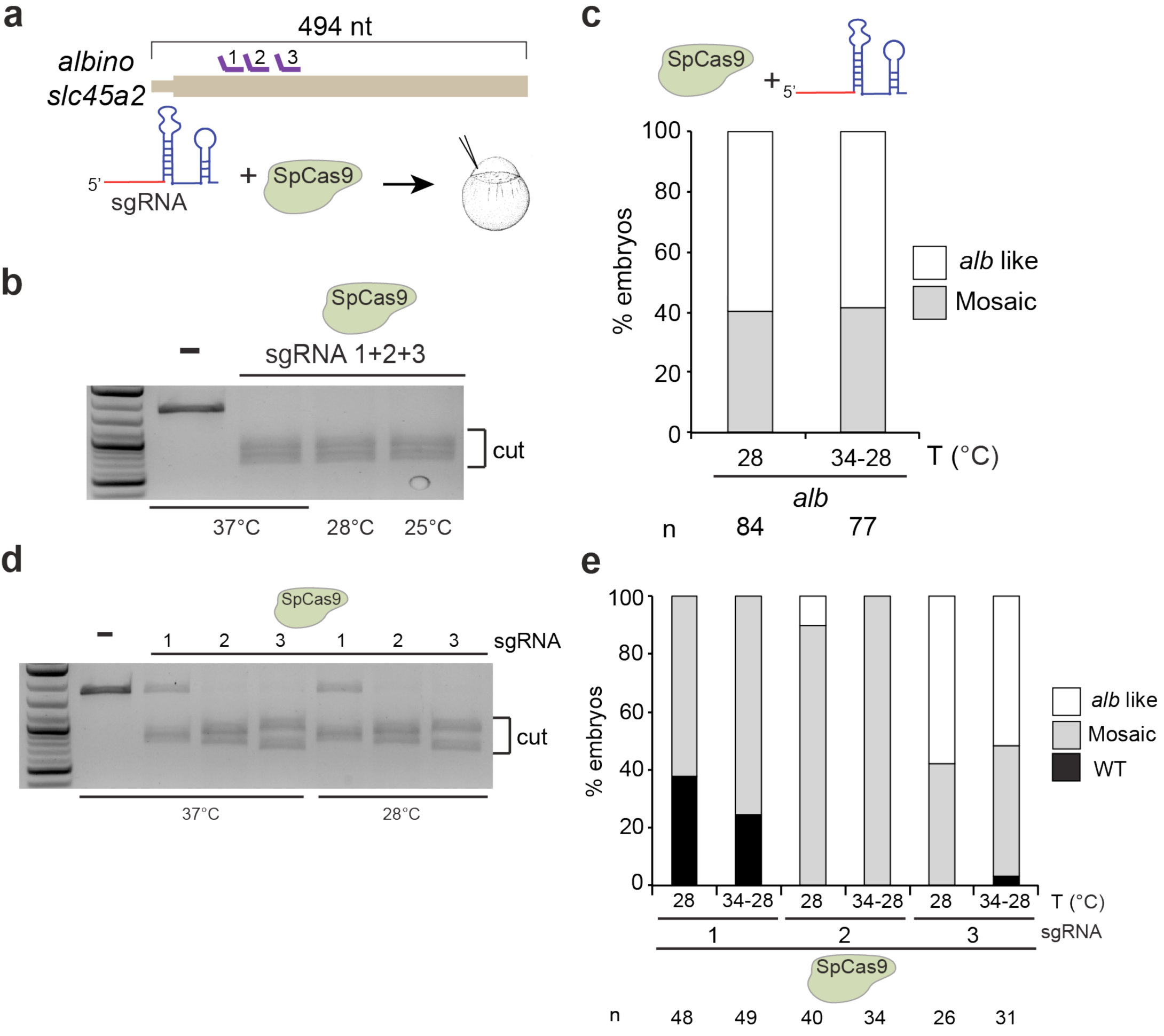
SpCas9 activity is not modulated by temperature in zebrafish. **a.** Diagram illustrating 3 sgRNAs (purple) targeting *slc45a2* exon 1 in zebrafish (top). Schematic showing sgRNAs (from above) that were assembled into SpCas9-sgRNA RNP complexes and injected into one-cell stage embryos (bottom). **b.** In vitro cleavage assay using SpCas9-sgRNA RNP complexes containing a mix of 3 sgRNAs (10 pmol) and a ~1.6 Kb (~ 0.11 pmol) PCR product containing sgRNA target sites for *slc45a2*. Incubations were carried out at 37°C, 28°C or 25°C for 90 min. **c.** Phenotypic evaluation of SpCas9-sgRNA RNP complexes (10 fmol) injections targeting *slc45a2* at different temperature incubations (T) as described in Fig. 2a. Stacked barplots showing the percentage of *alb*-like (white) and mosaic (gray) embryos 48 hpf after injection. χ^2^ test was performed and no significant differences were observed. **d.** *In vitro* cleavage assay using 10 pmol of SpCas9-sgRNA RNP complexes containing individual sgRNAs and a ~1.6 Kb (~ 0.11 pmol) PCR product including sgRNA target sites for *slc45a2*. Incubations were carried out at 37°C for 90 min. **e.** Phenotypic evaluation of individual SpCas9-sgRNA RNP complexes (10 fmol) injections targeting *slc45a2* at different temperature incubations (T) as described in Fig. 2a. Stacked barplots showing the percentage of *alb*-like (white), mosaic mutants (grey) and phenotypically WT (black) embryos 48 hpf after injection. Number of embryos evaluated (n) is shown for each condition. χ^2^ test was performed and no significant differences were observed.

**Supplementary Figure 7.**
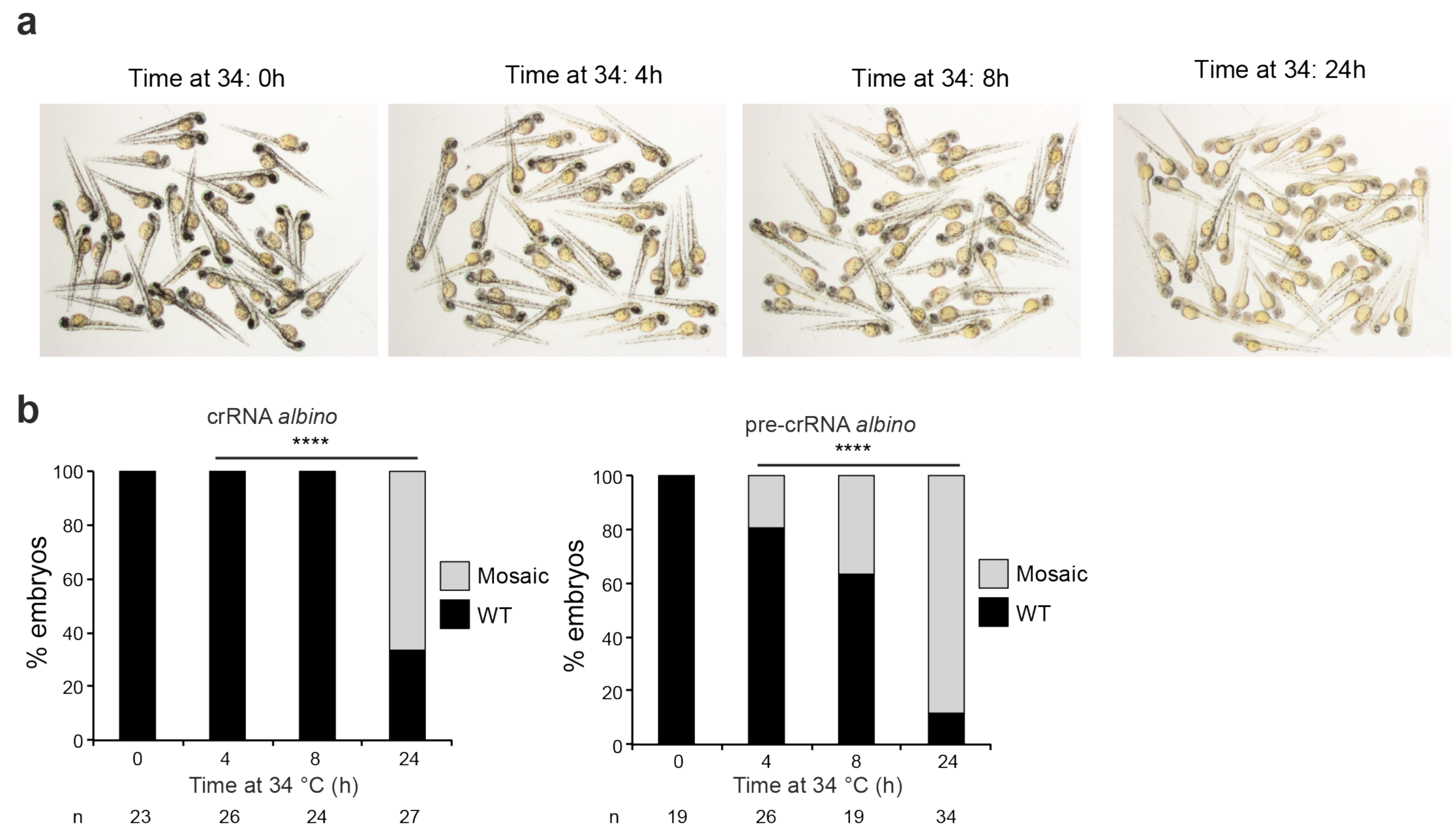
Longer incubations at 34°C improve AsCpf1 activity *in vivo.* **a.** A representative picture showing 48 hpf old embryos obtained after AsCpf1-crRNA RNP complexes injections targeting *tyr* in the conditions described in Fig. 2f. **b.** Phenotypic evaluation of crRNAs-AsCpf1 (left) or pre-crRNAs-AsCpf1 (right) RNP complexes (10 fmol) injections targeting *slc45a2* (*alb*) in the conditions described in Fig. 2f. Stacked barplots showing the percentage of mosaic mutants (grey), and phenotypically WT (black) embryos 48 hpf after injection. Number of embryos evaluated (n) is shown for each condition. χ^2^ test (****p< 0.0001).

**Supplementary Figure 8.**
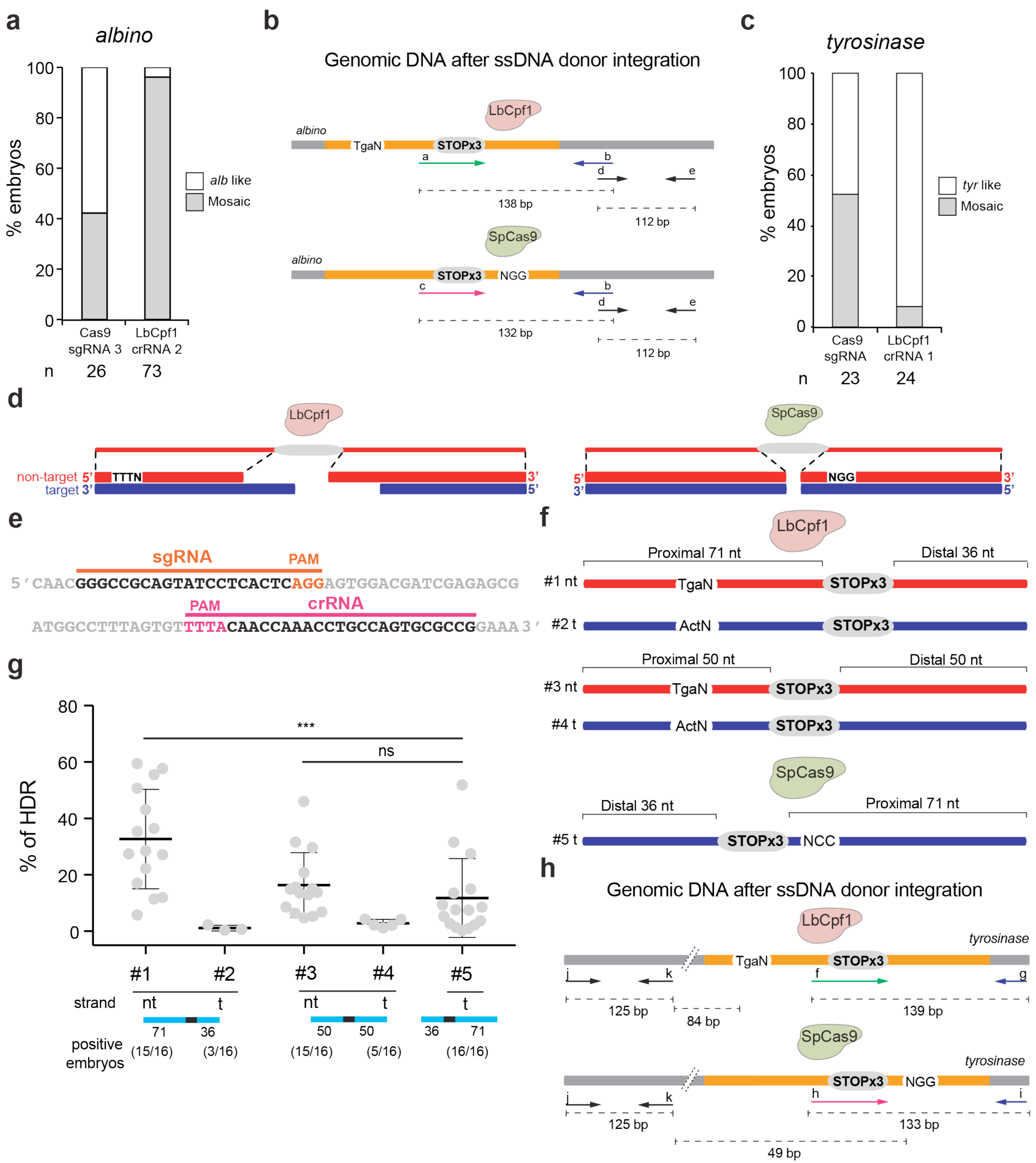
LbCpf1-mediated homology-directed repair. **a.** Phenotypic evaluation of Cas9-sgRNA 3 (Fig. 4b, Supplementary Fig. 6e) and LbCpf1-crRNA 2 (Fig. 4b, Supplementary Fig. 3f) RNP complexes (10 fmol) injections targeting *slc45a2* and used in Fig 4. Stacked barplots showing the percentage of *alb*-like (white), mosaic mutants (grey) embryos 48 hpf after injection. Number of embryos evaluated (n) is shown for each condition. **b.** Schema illustrating genomic DNA after ssDNA donor (containing 3 stop codons) integration in the *albino* locus. Specific primers (a,b,c) to amplify DNA integrations (orange) were used. Total DNA amount per embryo was calculated using primers (d,e) amplifying genomic DNA from the same locus (Supplementary Table 1, Methods). **c.** Phenotypic evaluation of Cas9-sgRNA and LbCpf1-crRNA 1 (Supplementary Table 1) RNP complexes (10 fmol) injections targeting *tyr and* used to quantify HDR. Stacked barplots showing the percentage of mosaic mutants (grey) and *tyr-*like (white) embryos 48 hpf after injection. Number of embryos evaluated (n) is shown for each condition. **d.** Schematic illustrating ssDNA donor centered in 3’ end of the LbCpf1-crRNA double-strand break (left) or in the blunt end of the SpCas9-sgRNA double-strand break (right). **e.** crRNA (pink line) and sgRNA (orange rline) target sequences in the *tyrosinase* locus used for this analysis. **f.** Schema illustrating different donor ssDNA (#1-#4) complementary to either the target strand (t) or non-target strand (nt) and with symmetric or asymmetric homology arms used in combination with LbCpf1-crRNA. PAM sequence was modified (TgaN/ActN) to prevent new editing post-HDR. An optimized ssDNA donor (#5) described for SpCas9-induced HDR^8^ was used in combination with SpCas9-sgRNA RNP as reference for comparison (bottom). **g.** qPCR quantification showing percentage of HDR from individual embryos when using LbCpf1 and different ssDNA donors in comparison with SpCas9. Results are shown as the averages ± standard errors of the means from 16 embryos in two independent experiments (n=8 embryos per experiment). Embryos showing a PCR amplification signal detectable by qPCR were considered positive. The data were analyzed by one-way ANOVA, followed by Bonferroni post-test for significance versus control condition (#5), (***) P < 0.001, (ns) not significant. **h.** Schema illustrating genomic DNA after ssDNA donor (containing 3 stop codons) integration into the *tyrosinase* locus. Specific primers (f-i) to amplify DNA integrations (orange) were used. Total DNA amount per embryo was calculated using primers (j,k) amplifying genomic DNA from the same locus (Supplementary Table 1, Methods).

